# Entry exclusion enables selective conjugative DNA delivery in synthetic bacterial communities

**DOI:** 10.64898/2026.09.24.754286

**Authors:** Kouhei Kishida, Natsumi Ogawa-Kishida, Leonardo Stari, Yoshiyuki Ohtsubo, Yuji Nagata

**Affiliations:** Department of Molecular and Chemical Life Sciences, Graduate School of Life Sciences, Tohoku University, 2-1-1 Katahira, Sendai 980-8577, Japan

## Abstract

Selective DNA delivery to specific members of assembled bacterial communities remains challenging. Bacterial conjugation enables efficient DNA delivery, but transfer to non-target recipients limits its specificity within mixed communities. Here, we repurpose plasmid entry exclusion (Eex) as a recipient-side gate to control conjugative DNA delivery. We demonstrate selective plasmid delivery to Eex-negative recipients within populations containing both Eex-expressing and Eex-negative cells. This recipient selectivity was maintained at increased cell densities and during prolonged mating. By combining RP4-type and F-type conjugation systems with their corresponding exclusion modules, we further directed DNA delivery from distinct donors to defined recipient populations. RP4-derived Eex also functioned in environmental bacteria, including *Pseudomonas putida* and *Sphingobium japonicum*, enabling recipient-specific exclusion within a multispecies mixture. Furthermore, repeated cycles of Eex-guided conjugation and selection altered community composition after assembly. These results establish entry exclusion as a recipient-side strategy for selective conjugative DNA delivery and, when combined with selection, for controlling the composition of assembled bacterial communities.

**GRAPHICAL ABSTRACT:** 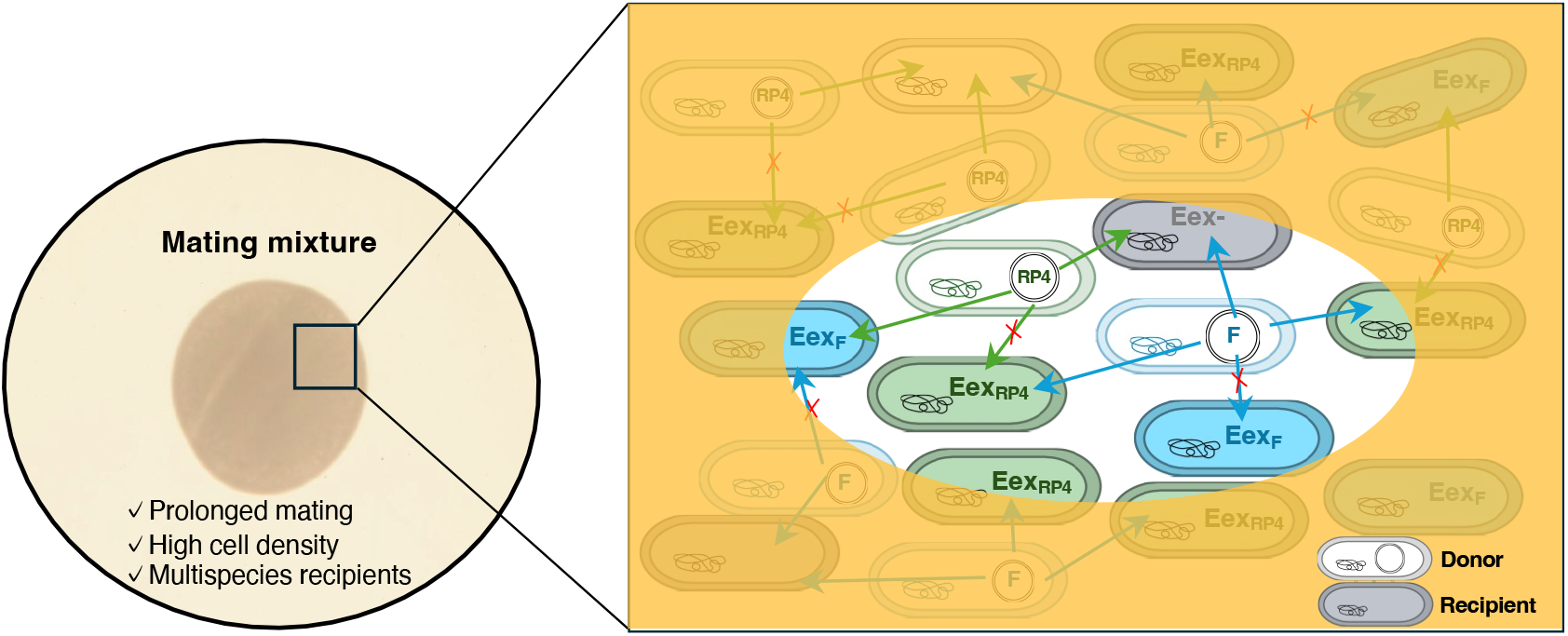

## INTRODUCTION

Bacteria form communities across diverse environments, including natural ecosystems and host-associated habitats, and support several key processes such as biogeochemical nutrient cycling, host nutrition and immune development, and protection against pathogen colonization (1–3). The complexity of natural communities often hinders controlled experimentation and causal inference, motivating the study of simplified synthetic bacterial communities (4,5). Defined synthetic bacterial communities are useful both for identifying principles that govern community dynamics and for engineering consortia with desired functions. However, methods for selectively delivering DNA to a specific member of an assembled multispecies synthetic bacterial community remain limited. This capability would allow researchers to genetically modify a specific strain within an established community and directly assess how that modification affects community behavior.

Bacterial conjugation is a practical and efficient method for DNA delivery into bacteria (6–8). Conjugation systems can transfer large DNA cargoes and achieve high transfer frequencies, making them useful for introducing genetic modules into bacterial cells. Some systems also have broad host ranges; for example, the conjugation system of the IncP-1 plasmid RP4 has been widely used to deliver DNA to diverse bacteria (9,10). These properties make conjugation attractive for genetic manipulation within microbial communities. However, transfer can occur to both target and non-target recipients, limiting the specificity of DNA delivery within mixed populations.

In Gram-negative bacteria, conjugative DNA transfer is initiated when a relaxase recognizes and nicks the origin of transfer (*oriT*), generating a transferable single-stranded DNA intermediate (11). This DNA is delivered from the donor to the recipient through a type IV secretion system (T4SS) (12). Although conjugation systems share a common overall mechanism for DNA processing and transport, their transfer machineries differ in composition and architecture. This variation may contribute to differences in transfer properties, including recipient range (13–15).

Many conjugative plasmids encode exclusion systems that limit transfer into recipients already harbouring related conjugative elements (16). These systems can act through surface exclusion, which interferes with mating-pair formation or stabilization, and entry exclusion (Eex), which blocks DNA transfer after donor–recipient contact has been established (16,17). Exclusion is often system-specific, with recipient-encoded factors suppressing transfer mediated by cognate conjugation machinery. By reducing recipient permissiveness to incoming DNA in a system-specific manner, Eex could serve as a recipient-side gate for selective DNA delivery within synthetic bacterial communities. To our knowledge, its use for directing plasmid transfer within mixed populations containing both Eex-expressing and Eex-negative recipients has not been experimentally demonstrated.

In this study, we developed an Eex-based strategy for selective DNA delivery within synthetic bacterial communities, using an RP4-type conjugation system as the primary transfer machinery. Recipient selectivity was confirmed at the single-cell level by fluorescence microscopy and was maintained at increased cell densities and during prolonged mating. Selectivity also extended to mobilizable plasmid transfer, including in triparental matings. By combining RP4- and F-type conjugation systems with their corresponding exclusion modules, we enabled parallel DNA delivery from distinct donors to defined recipient populations. RP4-derived Eex also supported selective transfer in environmental bacteria, including *Pseudomonas putida* KT2440 and *Sphingobium japonicum* UT26, and within mixed-species recipient populations. Finally, repeated cycles of Eex-guided conjugation and selection altered community composition. Together, these results establish Eex as a tool for selective DNA delivery and, when combined with selection, for manipulating the composition of assembled bacterial communities.

## MATERIAL AND METHODS

### Strains and growth conditions

*E. coli* strains listed in Table 1 were grown in Lysogeny Broth (LB) at 37 °C, and other species were grown in one-third-strength LB medium at 30 °C. LB contained (per liter) Bacto tryptone (10 g), Bacto yeast extract (5 g), and NaCl (5 g). One-third-strength LB contained (per liter) Bacto tryptone (3.3 g), Bacto yeast extract (1.7 g), and NaCl (5 g). For solid media, agar was added to a final concentration of 1.5% (w/v). For counterselection using *sacB*, NaCl was omitted from LB, and sucrose was added to a final concentration of 10% (w/v). Strains and plasmids were maintained with antibiotic selection as appropriate: ampicillin (100 μg/mL), tetracycline (20 μg/mL), kanamycin (50 μg/mL), nalidixic acid (20 μg/mL), streptomycin (50 μg/mL), gentamicin (10 μg/mL), chloramphenicol (50 μg/mL) and rifampicin (100 μg/mL). Because RHO3 is auxotrophic for diaminopimelic acid (DAP), DAP was added to the medium at a final concentration of 300 μg/mL during cultivation and mating of RHO3. Unless otherwise stated, expression from the P_BAD_ and P_lac_ promoters was induced by adding L-arabinose and IPTG to final concentrations of 10 mM and 1 mM, respectively.

**Table 1.**
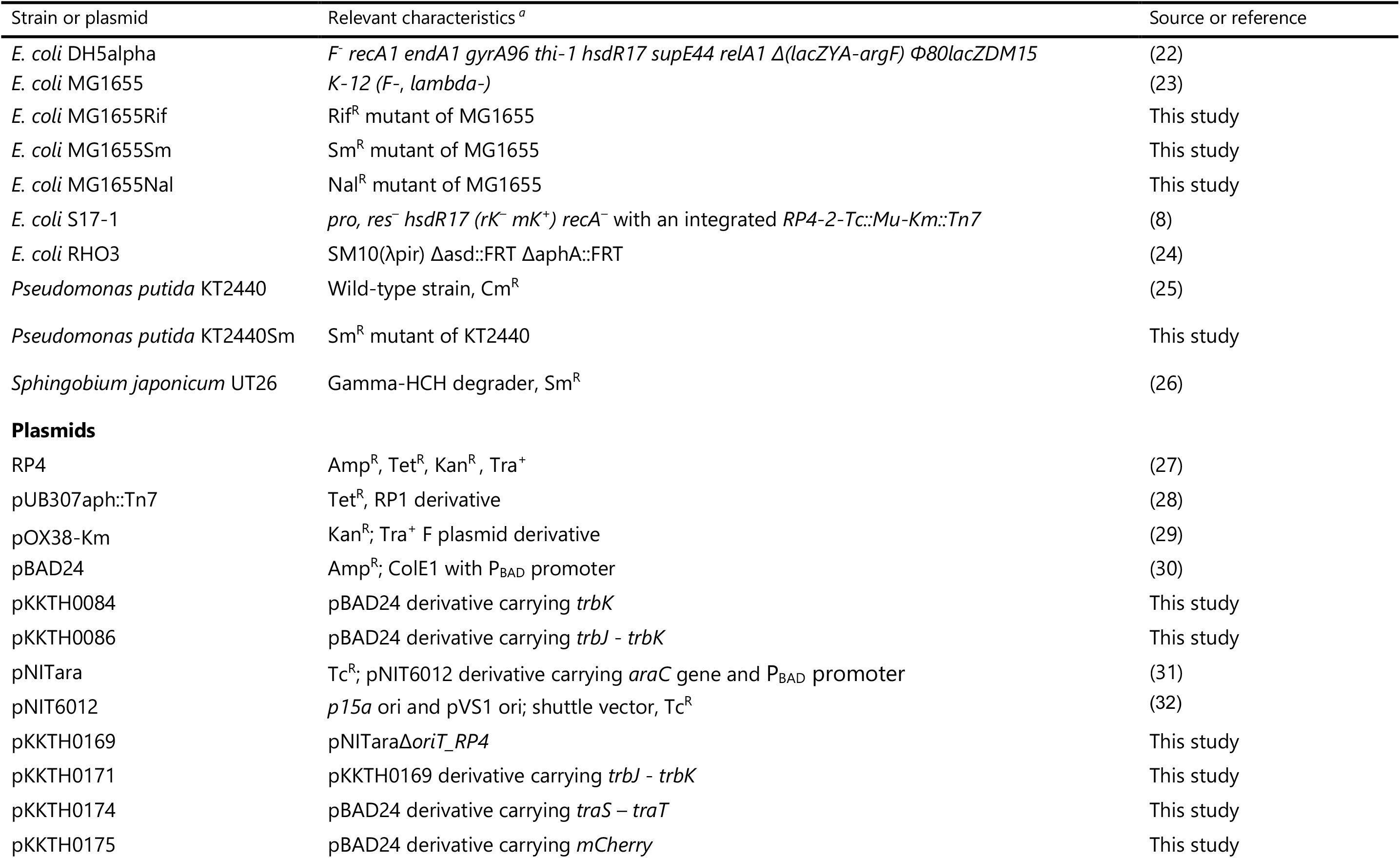
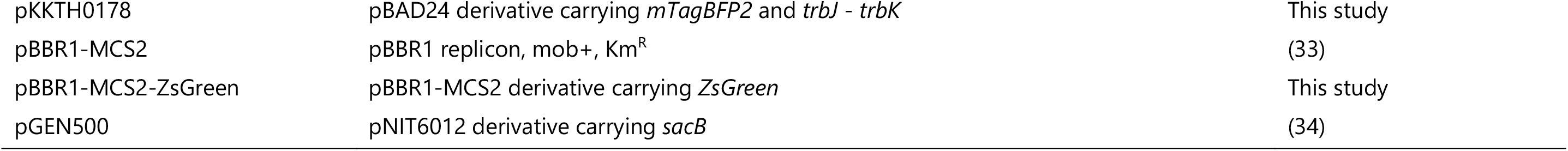
Bacterial strains and plasmids used in this study.

### Plasmid constructions

All plasmids and oligonucleotide primers used in these studies are listed in Tables 1 and S1, respectively. All plasmids are cloned by Gibson assembly kit. pKKTH0084, pKKTH0086, pKKTH0174 and pKKTH0175, each expressing a gene under P_BAD_ promoter, were constructed by amplifying individual gene regions and cloning them into the NheI site – HindIII site of pBAD24. pKKTH0178, expressing *trbJ* – *trbK* and *mTagBFP2* under P_BAD_ promoter, was constructed by amplifying *mTagBFP2* regions and cloning them into the HindIII of pKKTH0086. pKKTH0169, a derivative of pNITara lacking the RP4 origin of transfer (oriT_RP4), was generated by PCR amplification excluding oriT_RP4, followed by circularization of the resulting product. pKKTH0171, expressing *trbJ* – *trbK* under P_BAD_ promoter, was constructed by amplifying *trbJ* – *trbK* region and cloning it into the KpnI site of pKKTH0169. pBBR1-MCS2-ZsGreen expressing *ZSGreen* under P_lac_ promoter, was constructed by amplifying the fluorescent protein gene region and cloning them into the KpnI site – XhoI site of pBBR1-MCS2.

### Conjugation assays with *E. coli* recipients

Conjugation assays were performed using RP4, pUB307aph::Tn7, pOX38-Km, or the indicated mobilizable plasmids. RP4 and pUB307aph::Tn7 encode an RP4-type conjugation system but differ in their antibiotic resistance markers. Overnight cultures of donor and recipient strains were diluted 1:100 into fresh LB medium and incubated at 37 °C for a total of 2 h. For induction of Eex expression, L-arabinose was added to the recipient cultures after the first hour, and incubation was continued for another hour. Unless otherwise stated, this induction procedure was used for Eex-expressing strains.

Cells were collected by centrifugation and washed with LB. Donor and recipient suspensions were mixed at the indicated ratios. Unless otherwise stated, 250 μL of each strain suspension was used, yielding equal-volume mixtures. The mixtures were concentrated to 10 μL and spotted onto LB agar containing L-arabinose. After incubation at 37 °C for 30 min, cells were recovered from the agar surface in 1 mL LB, serially diluted, and plated on selective agar.

Transconjugants were selected using a plasmid-associated resistance marker together with a recipient-specific resistance marker. Recipient resistance to rifampicin, nalidixic acid, or streptomycin was used to counterselect donor cells. Kanamycin was used to select transconjugants carrying RP4 or pOX38-Km, whereas tetracycline was used to select those carrying pUB307aph::Tn7. Transfer frequencies were calculated as transconjugant CFU divided by donor CFU recovered after mating. For assays with unequal recipient mixing ratios (**Fig. 1D**), transfer frequencies were calculated separately for each recipient population as transconjugant CFU divided by the corresponding recipient CFU, to account for differences in recipient abundance.

**Figure 1.**
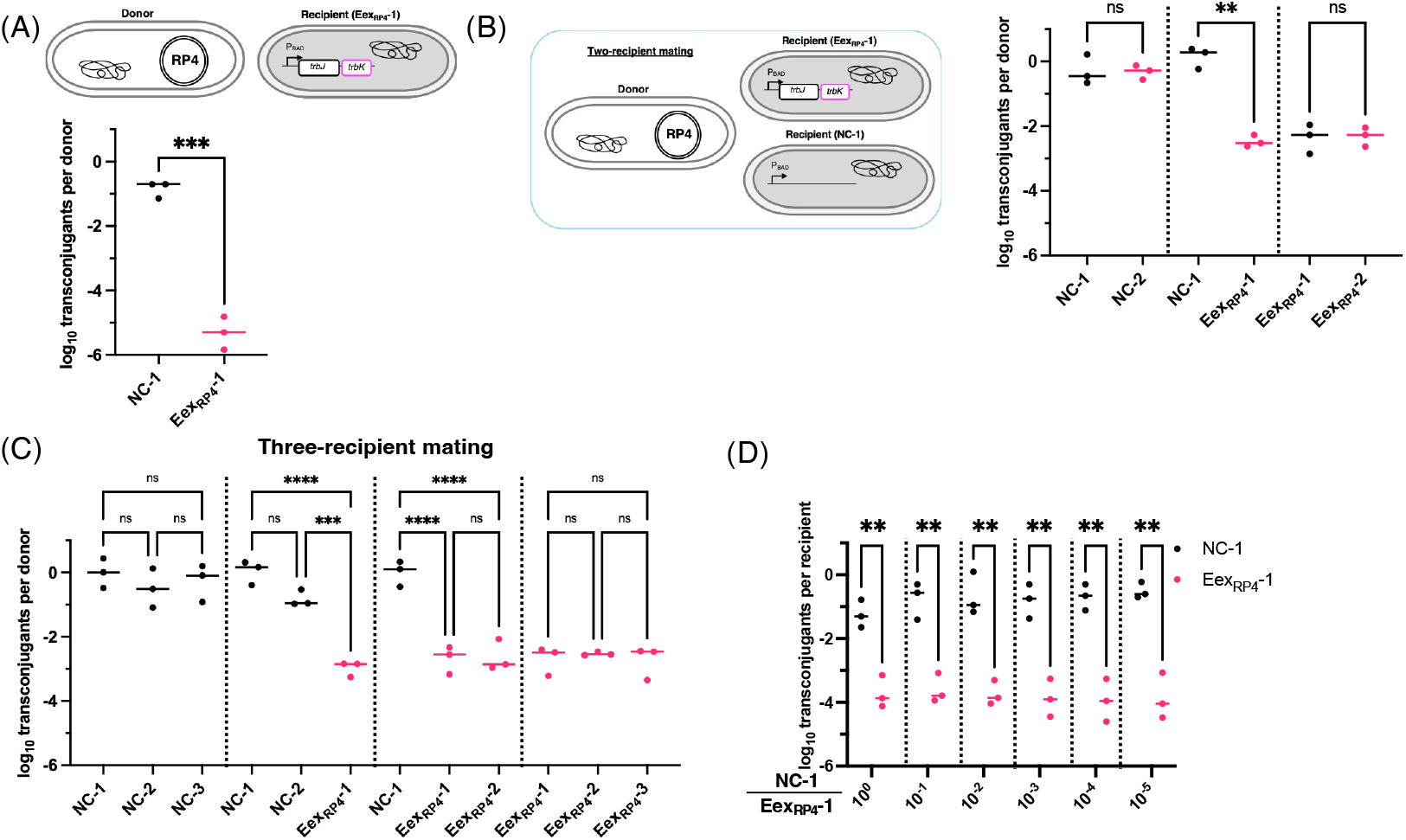
Entry exclusion selectively suppresses RP4-type conjugative transfer in mixed-recipient populations. **(A)** Schematic and quantification of conjugative transfer from a donor carrying an RP4-type plasmid into a control recipient (NC) and an Eex-expressing recipient (Eex_RP4_). The Eex_RP4_ recipient expressed the RP4 entry exclusion genes from the arabinose-inducible P_BAD_ promoter. Conjugation frequencies are expressed as log_10_ transconjugants per donor CFU. Statistical significance was determined by an unpaired two-tailed t-test; P = 0.0002.**(B)** Two-recipient mating assay. One donor carrying an RP4-type plasmid was mixed with two recipient strains in the same mating experiment. Recipient combinations included two NC recipients, one NC recipient and one Eex_RP4_ recipient, or two Eex_RP4_ recipients. Conjugation frequencies are expressed as log_10_ transconjugants per donor CFU. **(C)** Three-recipient mating assay. One donor carrying an RP4-type plasmid was mixed with three recipient strains in the same mating reaction. Recipient mixtures contained different combinations of NC and Eex_RP4_ recipients, as indicated on the x-axis. Conjugation frequencies are expressed as log_10_ transconjugants per donor CFU. **(D)** Selective transfer in recipient mixtures containing different ratios of NC and Eex_RP4_ recipients. NC-1 and Eex_RP4_-1 recipients were mixed at the indicated ratios and mated with the donor carrying an RP4-type plasmid. Conjugation frequencies are expressed as log_10_ transconjugants per recipient CFU. Black dots indicate NC recipients, and magenta dots indicate Eex_RP4_ recipients. Dots represent biological replicates, and horizontal bars indicate the mean. Statistical significance in panels B and C was assessed by ordinary one-way ANOVA followed by Tukey’s multiple-comparisons test on log₁₀-transformed transfer frequencies. In panel D, log₁₀-transformed transfer frequencies were compared using two-tailed paired t-tests, with values from the same mating mixture paired. Multiple testing across the six recipient mixing ratios was addressed using the two-stage step-up method of Benjamini, Krieger, and Yekutieli, with a false discovery rate of 1%. For panels A–C: ns, not significant; **P < 0.01; ***P < 0.001; ****P < 0.0001. For panel D: **P < 0.01.

For pNIT6012 transfer, *E. coli* S17-1 carrying pNIT6012 served as the donor, and transconjugants were selected with tetracycline and the appropriate recipient-specific antibiotic. For triparental transfer of pBBR1-MCS2, a helper strain carrying pUB307aph::Tn7, a cargo donor carrying pBBR1-MCS2, and the recipient strains were mixed as described above. Transconjugants carrying pBBR1-MCS2 were selected with kanamycin and the appropriate recipient-specific antibiotic.

### Conjugation assays with environmental bacterial recipients

The donor strain RHO3 carrying pBBR1-MCS2-ZsGreen was used to assess conjugative DNA delivery into *Pseudomonas putida* KT2440, *Sphingobium japonicum* UT26, and *E. coli* MG1655 derivatives. DAP was omitted from post-mating selective media to counterselect RHO3. All recipient strains in these assays were induced for 2 h with 10 mM L-arabinose in the presence of the appropriate antibiotics.

For assays with KT2440 recipients, 1 mL of an overnight culture grown at 30 °C in modified one-third-strength LB was added to 5 mL of fresh medium. L-arabinose and the appropriate antibiotics were added to their final working concentrations, and the culture was incubated at 30 °C for 2 h. One milliliter of the induced recipient culture was mixed with 1 mL of the RHO3 donor culture, and mating was performed on agar plates at 30 °C for 2 h.

For assays with UT26 recipients, cells were collected from agar plates and suspended in medium containing L-arabinose and the appropriate antibiotics. The suspension was incubated at 30 °C for 2 h. The induced recipient suspension and the RHO3 donor culture were mixed at 250 μL each, and mating was performed on agar plates at 30 °C for 2 h.

For mixed-recipient assays, KT2440, UT26, and rifampicin-resistant MG1655 derivatives were induced for 2 h with 10 mM L-arabinose. After induction, the three recipient suspensions were adjusted to the same OD₆₀₀ and mixed with the RHO3 donor suspension at a donor : KT2440 : UT26 : MG1655 volume ratio of 1:1:1:1. Mating was performed at 30 °C for 1.5 h. KT2440, UT26, and MG1655 derivatives were distinguished by selection with chloramphenicol, streptomycin, and rifampicin, respectively.

After mating, cells were recovered and plated on selective media as described above. For single-recipient assays with KT2440 or UT26 (**Fig. 5A, B**), transfer frequencies were calculated as transconjugant CFU divided by donor CFU recovered after mating. For mixed-recipient assays (**Fig. 5C**), transfer frequencies were calculated separately for each recipient population as transconjugant CFU divided by the corresponding recipient CFU recovered after mating. For the UT26 single-recipient assay, the detection limit was 10⁻⁶ transconjugants per donor, calculated from the donor CFU and the fraction of the recovered mating suspension plated for transconjugant enumeration. Samples yielding no transconjugant colonies were designated as not detected (N.D.) and plotted at the detection limit for visualization only. These samples were not treated as measured transfer frequencies for statistical testing.

### Detection of transconjugants by fluorescence microscopy

For microscopic detection of transconjugants, two-recipient mating assays were performed using a donor carrying a mobilizable ZsGreen-expressing plasmid and two differentially labeled recipient strains. The Eex-expressing recipient expressed mTagBFP2, whereas the control recipient expressed mCherry, both under the control of the P_BAD_ promoter. After mating for 1 h on LB agar supplemented with L-arabinose, cells were recovered from the agar surface, resuspended in LB medium, and mounted on glass slides. Transmitted-light and fluorescence images were acquired using an LSM 710 confocal laser-scanning microscope (Carl Zeiss) equipped with a 63× oil-immersion objective. ZsGreen, mTagBFP2, and mCherry were excited at 488, 405, and 561 nm, respectively. Five microscopic fields were analyzed for each of three independent biological replicates, with both recipient populations evaluated within the same fields.

Images were analyzed using Fiji/ImageJ. Recipient-cell masks were generated independently from the mCherry and mTagBFP2 channels using fixed fluorescence-intensity thresholds of 35–255 and 60–255, respectively, on an 8-bit intensity scale. Objects smaller than 0.4 µm² were excluded from the analysis. A binary ZsGreen-positive mask was generated using a fixed intensity threshold of 48–255 on the same scale. These threshold settings were established using a representative image and subsequently applied unchanged to all images. Candidate transconjugants were identified as recipient-cell regions for which at least 30% of the recipient-mask area overlapped with the ZsGreen-positive mask. Candidate double-positive cells were then visually inspected in the original fluorescence images to exclude apparent overlap caused by immediately adjacent ZsGreen-positive cells. A cell was classified as a transconjugant only when ZsGreen fluorescence colocalized with the corresponding recipient cell body. For each recipient population, the proportion of transconjugants was calculated for each biological replicate by dividing the total number of visually confirmed transconjugants across five microscopic fields by the total number of recipient cells across those fields.

### Sequential selection after conjugative plasmid transfer

Sequential conjugation–selection experiments were performed to examine how Eex-mediated plasmid exclusion affects community composition. RHO3 carrying pGEN500 served as the donor. The recipient community comprised three *E. coli* MG1655 derivatives: a streptomycin-resistant Eex-expressing strain and rifampicin- and nalidixic acid-resistant Eex-negative strains. Donor and recipient cultures were prepared as described above for conjugation assays with *E. coli* recipients. Aliquots of 250 μL from each culture were combined, concentrated by centrifugation, and spotted onto LB agar. Mating was performed at 37 °C for 1.5 h. For Eex induction, mating plates were supplemented with 10 mM L-arabinose.

After mating, cells were recovered from the agar surface and resuspended in LB. The recovered mixture was inoculated into either LB containing tetracycline or NaCl-free LB containing sucrose and incubated overnight at 37 °C. Following the initial conjugation–selection cycle, cultures underwent one additional cycle under tetracycline selection or two additional cycles under sucrose counterselection, for a total of two and three cycles, respectively. Each subsequent round included mating with the RHO3 donor carrying pGEN500, followed by overnight growth under the corresponding selection condition. At each passage, 5 μL was inoculated into 5 mL of fresh selection medium. DAP was omitted from selection cultures and enumeration plates to counterselect RHO3.

Population composition was determined by selective plating after the initial mating and after each round of selection. The three recipient populations were enumerated separately on plates containing streptomycin, rifampicin, or nalidixic acid. The relative abundance of each recipient population was calculated by dividing its CFU by the sum of the CFU of all three recipient populations.

### Statistical analysis

All statistical analyses were performed using GraphPad Prism 9. The statistical tests, data transformations, pairing of observations, and procedures used to account for multiple testing are specified in the corresponding figure legends. Statistical significance is indicated in the figures.

## RESULTS

### Recipient-mediated entry exclusion enables selective plasmid transfer in mixed populations

To establish a selective conjugation system based on entry exclusion, we employed the broad-host-range plasmid RP4 in *E. coli*. Conjugation assays were performed using solid-surface mating conditions, under which RP4 transfer is highly efficient. Although *trbK* is annotated as the entry exclusion (Eex) determinant of RP4, previous studies have suggested that functional exclusion requires both *trbK* and the adjacent gene *trbJ* (18). Consistent with this notion, recipient cells harboring a plasmid expressing trbK alone showed only a modest reduction in transfer frequency, approximately 3.5-fold relative to the empty-vector control (**Fig. S1**). We therefore constructed a recipient strain (Eex_RP4_-1) carrying a plasmid expressing both *trbJ* and *trbK* under the control of the arabinose-inducible P_BAD_ promoter. Induced expression of *trbJ* – *trbK* did not detectably affect growth under the tested conditions (**Fig. S2(A)**). Under inducing conditions, RP4 transfer into Eex_RP4_-1 was reduced by approximately 10^4^-fold relative to the empty-vector control strain (NC-1) (**Fig. 1A**). In contrast, no significant reduction in transfer frequency was observed in the absence of induction, indicating tight regulatory control of exclusion activity (**Fig. S3**). To determine whether entry exclusion enables selective transfer in mixed populations (**Fig. 1B**), we performed two-recipient mating assays using pairs of recipient strains carrying distinct antibiotic resistance markers. In each assay, both recipients were exposed to the same donor population, and transconjugants from each recipient population were enumerated separately. When NC-1 and Eex_RP4_-1 were combined, transfer into NC-1 occurred at a substantially higher frequency than into Eex_RP4_-1, demonstrating selective plasmid delivery within the same mating mixture. When both recipients carried empty vectors (NC-1 and NC-2), transfer frequencies were approximately 10⁻¹ for both strains. Conversely, when both recipients expressed *trbJ*–*trbK* (Eex_RP4_-1 and Eex_RP4_-2), transfer frequencies were reduced to approximately 10⁻^2^ for both strains. We next extended this analysis to a three-recipient mating system (**Fig. 1C**). Selective plasmid transfer was similarly observed, enabling preferential delivery to either two of the three recipients or to a single designated recipient, depending on the combination of Eex-expressing strains present. These results indicate that entry exclusion can be harnessed to direct plasmid flow within more complex recipient communities.

To examine whether selective transfer depends on recipient composition, we systematically varied the ratio of NC-1 to Eex_RP4_-1 (NC-1:Eex_RP4_-1 =1:1 to 10⁻⁵) in two-recipient mating assays (**Fig. 1D**). For these assays, transfer frequencies were calculated as transconjugant CFU divided by the corresponding recipient CFU recovered after mating. Across all mixing ratios, transfer into NC-1 remained significantly higher than into EexRP4-1.

Even when NC-1 represented a minor fraction of the population (NC-1:EexRP4-1 = 1:10⁵), its transfer frequency remained approximately two orders of magnitude higher than that of Eex_RP4_-1. Thus, recipient selectivity was maintained even when Eex-negative recipients were greatly outnumbered by Eex-expressing recipients. Collectively, these findings demonstrate that inducible co-expression of *trbJ* and *trbK* confers RP4 entry exclusion and enables selective plasmid delivery within mixed-recipient populations across the tested recipient ratios.

### Expansion of entry-exclusion-based selective conjugation to mobilizable plasmids and triparental mating

To test whether entry-exclusion-based selectivity applies to mobilizable plasmid transfer, we first used a triparental mating setup in which RP4 provided the transfer machinery in trans (**Fig. S4A**). In this setup, a helper donor carrying RP4-type plasmid, a second donor carrying the mobilizable plasmid pBBR1 plasmid, and two recipient strains (NC-1 and Eex_RP4_-1) were mixed. Transfer of pBBR1 plasmid into NC-1 was readily detected, whereas transfer into Eex_RP4_-1 was strongly reduced (∼10⁴-fold), indicating that recipient entry exclusion effectively blocks mobilizable plasmid transfer driven by RP4.

We next examined selective transfer in a standard donor–recipient mating using *E. coli* S17-1 carrying pNIT6012, which contains *oriT*_RP4_ (**Fig. S4B**). Consistent with the triparental assay, conjugation into NC-1 was significantly higher than into Eex_RP4_-1, confirming that entry exclusion confers selective suppression of *oriT_RP4_*-dependent DNA transfer. Together, these results show that entry-exclusion-mediated selectivity is not limited to self-transmissible RP4 transfer but also extends to mobilizable plasmids and *oriT*_RP4_-dependent cargo transfer, enabling controlled DNA flow in mixed populations.

### Selective plasmid transfer is observed at the single-cell level

To directly visualize selective plasmid transfer, we performed two-recipient mating followed by fluorescence microscopy (**Fig. 2A**). The donor strain carried an RP4-type plasmid pUB307aph::Tn7 and a mobilizable plasmid encoding ZsGreen, while recipient strains were labeled with mCherry (NC-1) or mTagBFP2 (Eex_RP4_-1). Colony-based conjugation assays confirmed that recipient selectivity was maintained in the fluorescently labeled strains (**Fig. 2B**). Comparison of the two recipient populations within the same microscopic fields revealed that ZsGreen-positive cells were predominantly found among mCherry-positive NC-1 recipients (**Fig. 2C**). Quantitative image analysis followed by visual verification identified 1,916 transconjugants among 3,466 NC-1 cells (55.3%), compared with only one transconjugant among 5,688 Eex_RP4_-1 cells (0.0176%), pooled across three independent biological replicates (**Fig. 2D**). The proportions of transconjugants ranged from 51.3% to 59.4% in NC-1 and from 0% to 0.044% in Eex_RP4_-1 across the biological replicates. These findings confirm that entry exclusion enables selective plasmid delivery to Eex-negative recipients within mixed populations at single-cell resolution.

**Figure 2.**
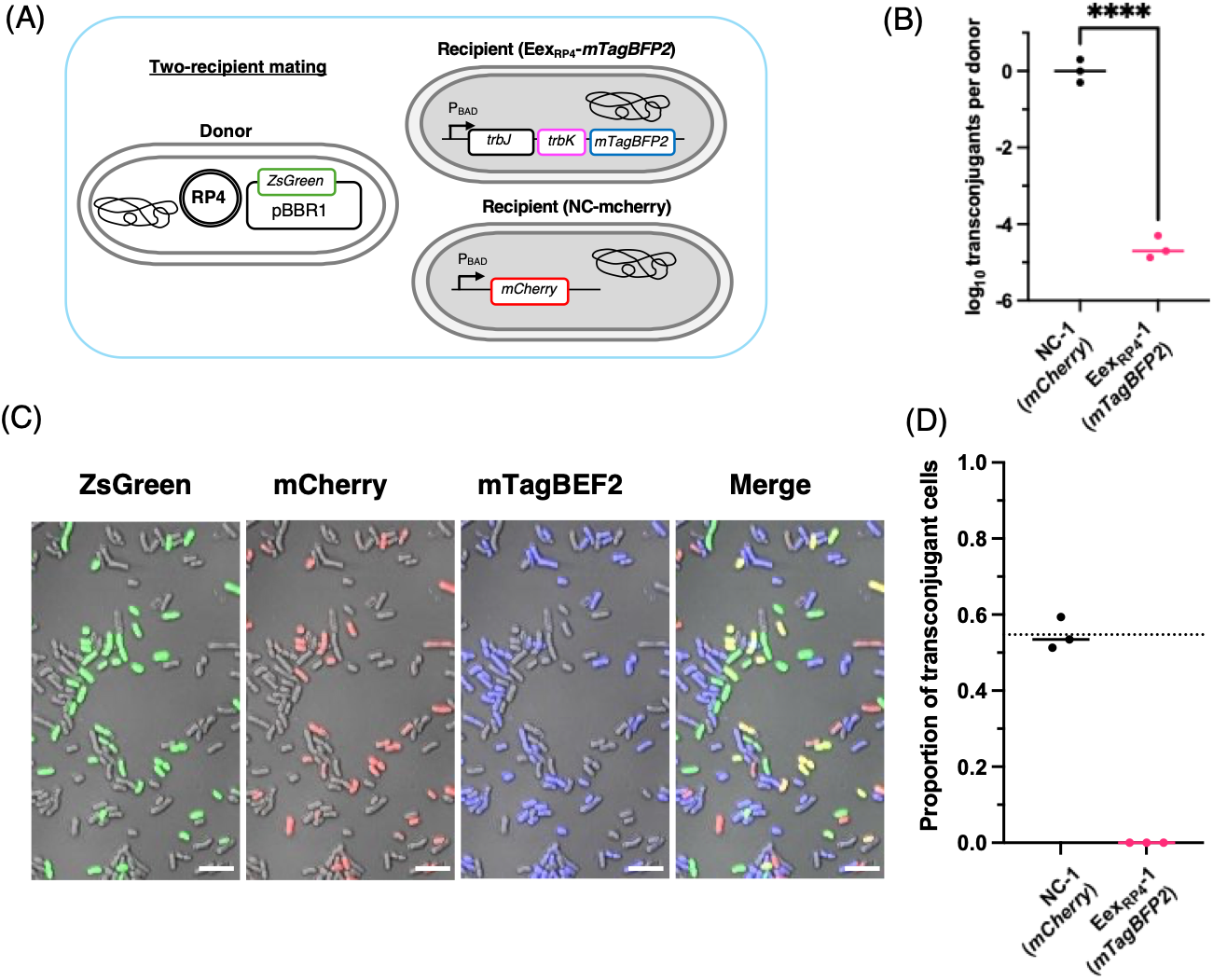
Single-cell fluorescence microscopy confirms selective plasmid acquisition in mixed-recipient mating. **(A)** Schematic of the two-recipient mating assay used for fluorescence microscopy. The donor carried an RP4-type conjugative plasmid and the mobilizable ZsGreen-expressing plasmid pBBR1-MCS2-ZsGreen. The Eex_RP4_-1 recipient was labeled with mTagBFP2, and the control recipient NC-1 was labeled with mCherry. **(B)** Quantification of conjugative transfer by a colony-based assay using the same donor and recipient combination. Transfer frequencies were calculated as transconjugant CFU per donor CFU recovered after mating and are shown on a log₁₀ scale. Statistical significance was determined using an unpaired two-tailed t-test on log₁₀-transformed values. ****P < 0.0001. **(C)** Representative transmitted-light and fluorescence images after mating. ZsGreen marks donor cells and recipients that acquired the mobilizable plasmid; mCherry and mTagBFP2 mark NC-1 and Eex_RP4_-1 recipients, respectively. The merged image shows an overlay of the three fluorescence channels and the transmitted-light image. Scale bars, 5 μm. **(D)** Proportion of transconjugant cells within each recipient population. Transconjugants were identified by image analysis followed by visual verification, as described in Materials and Methods. Five fields were analyzed for each of three independent biological replicates. For each recipient population, the proportion was calculated by dividing the total number of confirmed transconjugants across the five fields by the total number of corresponding recipient cells. In panels B and D, black and magenta dots indicate NC-1 and Eex_RP4_-1 recipients, respectively. Each dot represents one biological replicate (n = 3), and horizontal bars indicate the mean.

### Selective plasmid transfer is maintained during prolonged mating and at increased cell densities

Prolonged mating and increased cell density could compromise recipient selectivity by increasing opportunities for transfer into non-target cells. Having confirmed selective plasmid acquisition at the single-cell level, we therefore examined whether Eex-mediated selectivity was maintained under these conditions. We first varied the mating duration from 30 min to 96 h using a donor carrying RP4 and a mixed population of NC-1 and Eex_RP4_-1 recipients. Transfer into Eex_RP4_-1 remained significantly lower than into NC-1 at all tested mating durations (**Fig. 3A**). Although transfer into Eex_RP4_-1 increased with extended mating, it remained approximately three orders of magnitude lower than into NC-1 after 24 h. We next varied the cell density over a 100-fold range by increasing the total number of donor and recipient cells from 10⁷ to 10⁹ per mating mixture while keeping the mating volume and surface area constant. After 30 min of mating, transfer into Eex_RP4_-1 remained significantly lower than into NC-1 at all tested cell densities (**Fig. 3B**). Together, these results demonstrate that Eex-mediated selectivity is maintained within mixed-recipient populations during prolonged mating and at increased cell densities.

**Figure 3.**
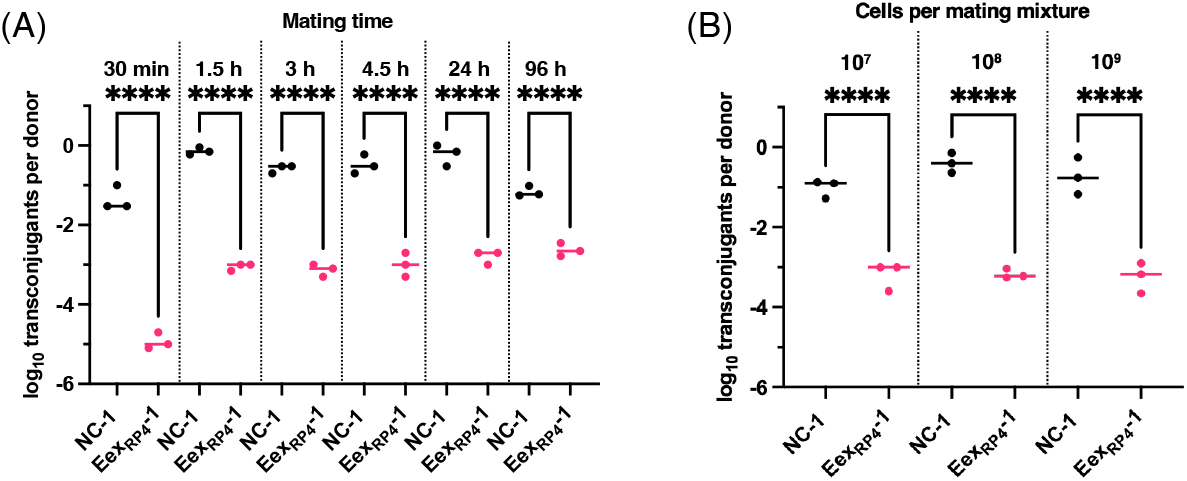
RP4-mediated entry exclusion is maintained across different mating durations and total cell numbers. (A) Conjugative transfer from a donor carrying RP4 to a mixed-recipient population containing NC-1 and Eex_RP4_-1 after mating for 30 min, 1.5 h, 3 h, 4.5 h, 24 h, 96 h. (B) Conjugative transfer using the same donor– recipient combination after 30 min of mating with 10⁷, 10⁸, or 10⁹ total cells per mating mixture, including both donors and recipients. Transfer frequencies are expressed as log₁₀ transconjugants per donor. Each dot represents an independent biological replicate (n = 3), and horizontal lines indicate means. Statistical significance was assessed using ordinary one-way ANOVA followed by Tukey’s multiple-comparison test on log₁₀-transformed transfer frequencies. ****P < 0.0001.

### Plasmid-specific exclusion directs distinct plasmids to defined recipient populations

Entry exclusion systems are generally specific to cognate or closely related conjugation systems (17,19). To test whether this specificity could be exploited to program dual-plasmid delivery, we combined the RP4 system with the F plasmid conjugation machinery. We constructed a recipient strain, designated EexF-1, expressing the F plasmid entry exclusion factor TraS and surface exclusion factor TraT under the control of the arabinose-inducible P_BAD_ promoter. Because high-level induction impaired growth (**Fig. S2B**), 0.01 mM L-arabinose was used for assays involving TraS–TraT expression. In three-recipient mating assays (**Fig. 4A**), recipients expressing TraS–TraT strongly suppressed pOX38-Km acquisition while remaining permissive to transfer of the RP4-type plasmid (**Fig. 4B**). Conversely, recipients expressing RP4-derived Eex strongly suppressed acquisition of the RP4-type plasmid while remaining permissive to pOX38-Km transfer. Although transfer frequencies into recipients expressing the non-cognate exclusion system differed from those into NC-1, each plasmid was transferred more efficiently into recipients expressing the non-cognate rather than the cognate exclusion system.

**Figure 4.**
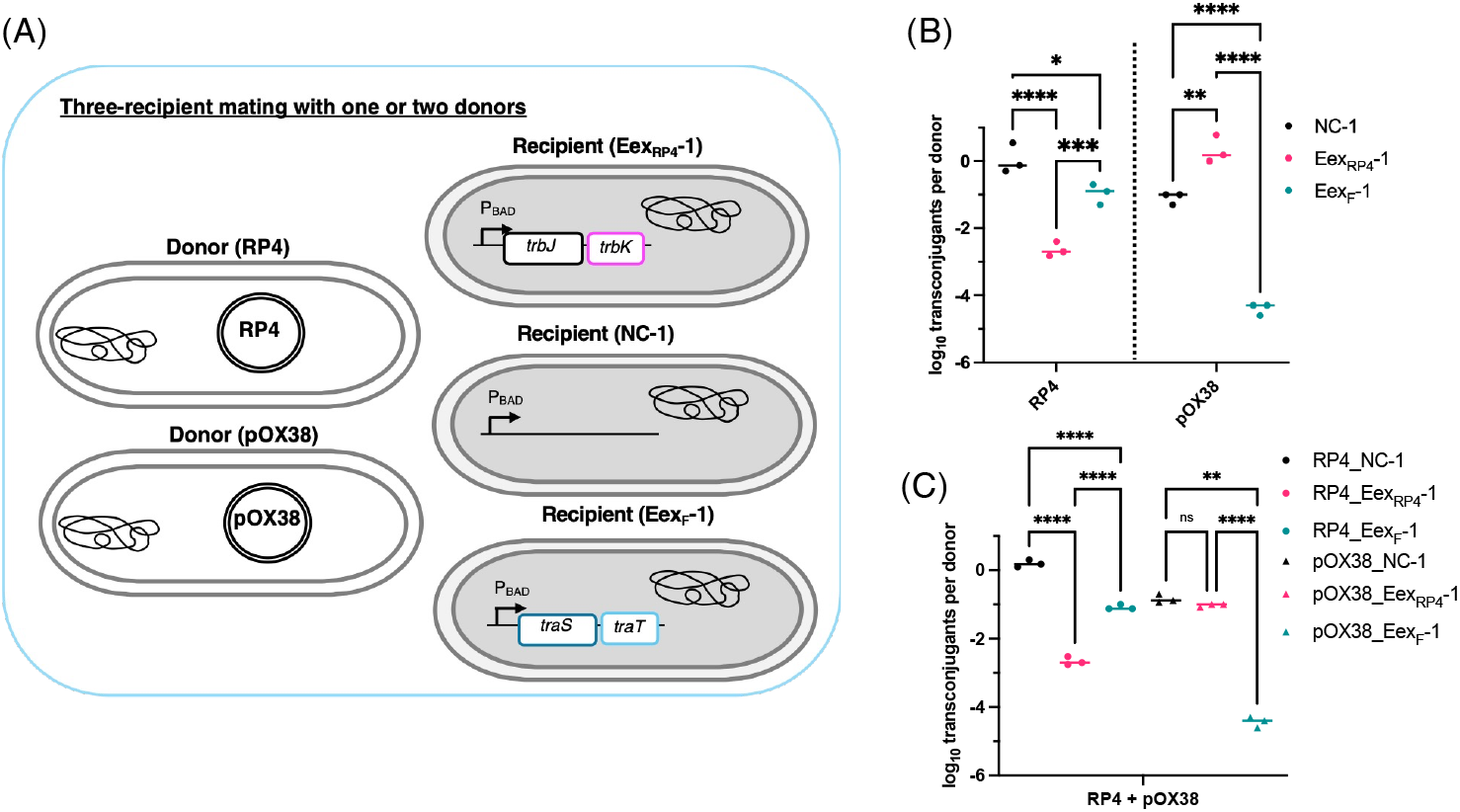
Orthogonal entry exclusion systems enable selective routing of conjugative DNA delivery from distinct donors. **(A)** Schematic of three-recipient mating with single or dual donor input. Donors carried either an RP4-type plasmid (pUB307aph::Tn7) or the F-derived conjugative plasmid pOX38-Km. Recipient mixtures contained a control recipient, an Eex_RP4_ recipient expressing the RP4 entry exclusion genes *trbJ* and *trbK*, and an Eex_F_ recipient expressing the F plasmid entry exclusion gene *traS* and surface exclusion gene *traT*. **(B)** Selective conjugative transfer from a single donor. The pUB307aph::Tn7 donor or the pOX38-Km donor was individually mixed with the three-recipient population. Transfer from the pUB307aph::Tn7 donor was selectively reduced in the Eex_RP4_ recipient, whereas transfer from the pOX38-Km donor was selectively reduced in the Eex_F_ recipient. **(C)** Selective conjugative transfer from dual donors. pUB307aph::Tn7 and pOX38-Km donors were mixed together with the three-recipient population in the same mating experiment. Transfer from each donor was selectively reduced in the recipient expressing the corresponding Eex module. Conjugation frequencies were calculated as transconjugants per donor CFU and are shown on a log_10_ scale. Dots represent biological replicates, and horizontal bars indicate the mean. Statistical significance in panel B was assessed by two-way ANOVA followed by Šídák’s multiple-comparisons test. In panel C, log₁₀-transformed transfer frequencies were analyzed by repeated-measures one-way ANOVA with Geisser–Greenhouse correction, followed by Tukey’s multiple-comparisons test across all 15 pairwise comparisons. Measurements from the same mating mixture were matched. ns, not significant; *P < 0.05; **P < 0.01; ***P < 0.001; ****P < 0.0001. P values for pairwise comparisons were adjusted for multiple testing.

To determine whether this specificity could direct simultaneous delivery from two donor populations, we performed mixed-donor mating assays in which a donor carrying an RP4-type plasmid and a donor carrying pOX38-Km were combined with the recipient populations in a single mating mixture. Under these conditions, each plasmid was preferentially delivered to recipients lacking its cognate exclusion system, demonstrating that recipient selectivity was maintained when both donor populations were present (**Fig. 4C**). Together, these results demonstrate that distinct exclusion systems can be combined to direct different conjugative plasmids to defined recipient populations within the same mating mixture.

### Entry-exclusion-based selectivity extends to environmental bacterial species

To determine whether RP4-derived entry exclusion functions beyond *E. coli*, we examined two environmental bacterial species, *Pseudomonas putida* KT2440 and *Sphingobium japonicum* UT26. The *trbJ–trbK* genes were cloned into the broad-host-range plasmid pNITara under the control of an arabinose-inducible promoter. Upon induction, Eex expression reduced conjugative transfer into KT2440 by approximately three orders of magnitude relative to the control strain **(Fig. 5A**). In UT26, no transconjugants were detected in any of the three independent biological replicates of the Eex-expressing recipient, with a detection limit of 10⁻⁶ transconjugants per donor CFU (**Fig. 5B**). These results demonstrate that RP4-derived Eex functions in both environmental bacterial hosts.

**Figure 5.**
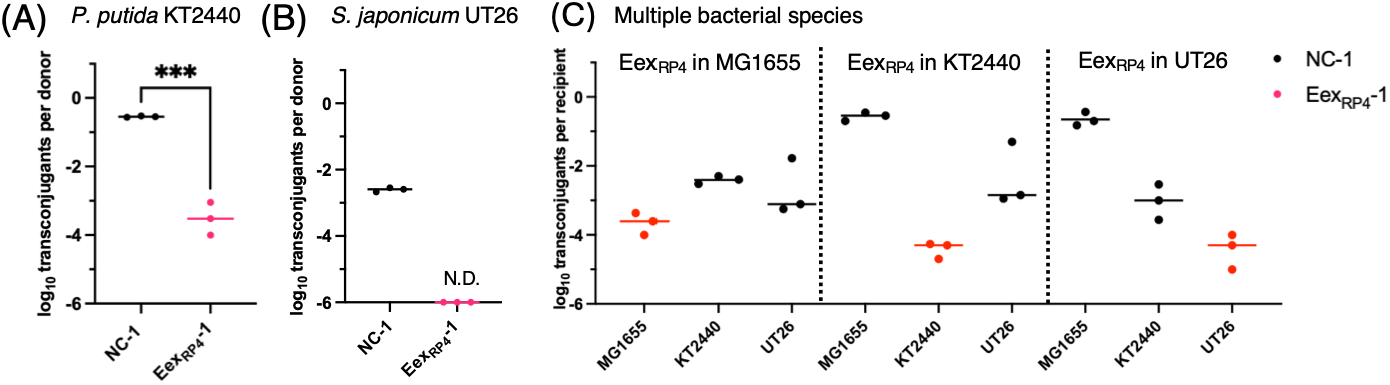
RP4-derived entry exclusion functions in environmental bacterial recipients and multi-species recipient mixtures. (**A, B**) Conjugative transfer into control (NC-1) and Eex-expressing (Eex_RP4_-1) recipients of *Pseudomonas putida* KT2440 (A) and *Sphingobium japonicum* UT26 (B). Eex-expressing recipients carried *trbJ* and *trbK* under the control of an arabinose-inducible promoter on pNITara. Transfer frequencies were calculated as transconjugants per donor CFU and plotted on a log₁₀ scale. In panel B, no transconjugants were detected in any of the three biological replicates of the Eex-expressing recipient. These observations are plotted at the detection limit of 10⁻⁶ transconjugants per donor CFU for visualization only. N.D., not detected. (**C**) Mixed-species mating assays with *E. coli* MG1655, *P. putida* KT2440, and *S. japonicum* UT26 as recipients. In each mating mixture, the RP4 exclusion module was expressed in only one recipient species, as indicated above each group. Recipient populations and their transconjugants were enumerated by selective plating. Transfer frequencies were calculated as transconjugants per CFU of the corresponding recipient species and plotted on a log₁₀ scale. Black dots indicate control recipients, and colored dots indicate Eex-expressing recipients. Each dot represents an independent biological replicate (n = 3). Horizontal lines indicate means of the log₁₀-transformed frequencies, except for the undetected group in panel B. Statistical significance in panel A was assessed using an unpaired two-tailed t-test on log₁₀-transformed transfer frequencies. ***P < 0.001. No statistical comparison was performed for panel B.

We next asked whether Eex-based selectivity could operate in a mixed-recipient population containing multiple bacterial species (**Fig. 5C**). For this experiment, *E. coli* MG1655, KT2440, and UT26 were mixed as recipients, and RP4-derived Eex was expressed in one recipient species at a time. In each combination, transfer into the Eex-expressing population was lower than into the same species in mixtures where it lacked Eex, while the other two recipient species remained permissive to plasmid acquisition. These results show that RP4-derived Eex can impose recipient-specific exclusion within a multispecies recipient population, extending its application to selective DNA delivery in synthetic communities composed of different bacterial species.

### Sequential conjugation and selection alter recipient population composition

We next investigated whether entry exclusion-based selective conjugation could be used to control the composition of recipient populations. To this end, we used pGEN500, a plasmid carrying a tetracycline resistance marker and the counterselectable marker *sacB*, together with an RP4 transfer-proficient donor. Cells that acquired pGEN500 could be enriched by tetracycline selection, whereas sucrose counterselection could favor cells that had avoided plasmid acquisition (**Fig. 6A**). A mixed recipient population consisting of Eex_RP4_-1, NC-1, and NC-2 was subjected to successive cycles of conjugation and selection. Under tetracycline selection, the relative abundance of Eex _RP4_-1 decreased markedly, enriching Eex-negative recipients. In the representative experiment shown, NC-1 became the dominant population by the final sampling point (**Fig. 6B**). This result is consistent with reduced acquisition of pGEN500 by EexRP4-1 due to entry exclusion. In contrast, successive cycles of conjugation and sucrose counterselection progressively enriched Eex _RP4_-1, which comprised nearly the entire recipient population at the final sampling point in the representative experiment shown. This enrichment is consistent with reduced acquisition of the *sacB*-carrying plasmid by Eex _RP4_-1, allowing these recipients to escape sucrose counterselection. Together, these findings demonstrate that Eex-mediated differences in plasmid acquisition can be coupled to selection to shift the composition of an assembled bacterial population. Tetracycline selection favored Eex-negative recipients, whereas sucrose counterselection favored Eex-expressing recipients.

**Figure 6.**
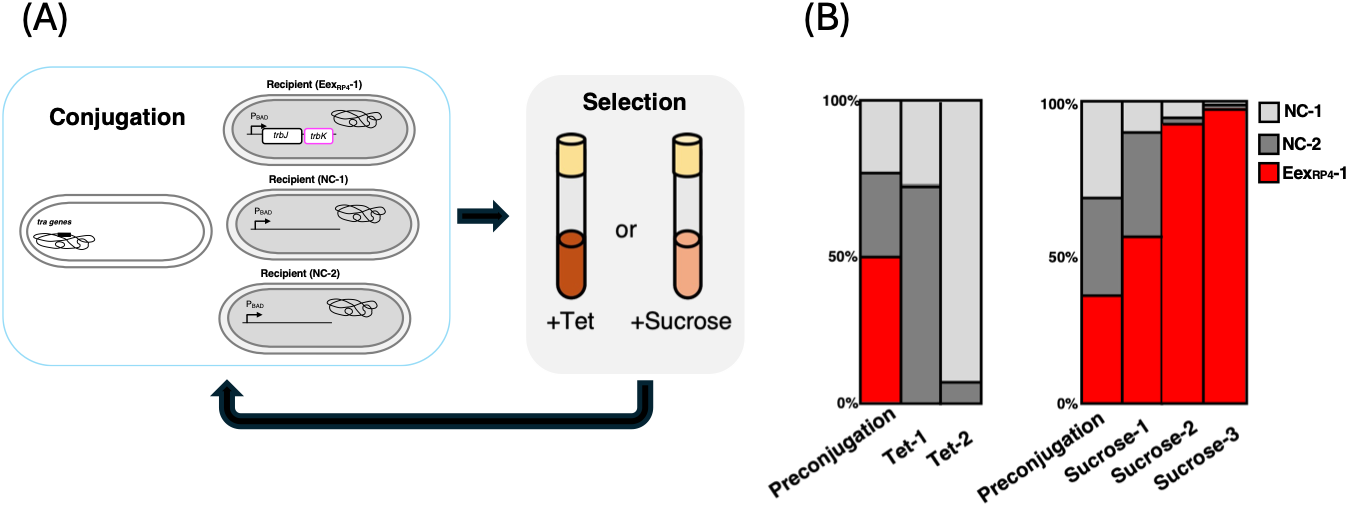
**(A)** Experimental design. An RP4 transfer-proficient donor carrying pGEN500, which confers tetracycline resistance and carries *sacB*, was mixed with a recipient community consisting of an RP4 entry exclusion-expressing strain (Eex_RP4_-1) and two non-exclusion control strains (NC-1 and NC-2). **(B)** Relative abundance of Eex _RP4_-1, NC-1, and NC-2 before conjugation and after successive cycles of conjugation and tetracycline selection or sucrose counterselection. Numbers indicate the number of conjugation–selection cycles completed. Stacked bars show the relative abundance of each recipient population in one representative experiment. Similar trends in Eex _RP4_-1 depletion or enrichment were observed in three independent biological replicates.

## DISCUSSION

In this study, we repurposed entry exclusion as a recipient-side gate for selective conjugative DNA delivery in synthetic bacterial communities. Entry exclusion has traditionally been understood as a plasmid-encoded mechanism that limits redundant acquisition of related conjugative elements (8,16). To our knowledge, its use to direct selective plasmid transfer within a mixed population containing both Eex-positive and Eex-negative recipients has not previously been experimentally demonstrated. By exposing both recipient populations to the same donors in a single mating mixture, we showed that Eex selectively suppressed transfer into Eex-expressing cells while Eex-negative cells remained permissive (**Figs. 1 and 2**). These findings establish that Eex can be used to determine the recipients of conjugative DNA within assembled bacterial communities.

Previous approaches to targeted conjugation have mainly focused on increasing donor-recipient contact. Synthetic adhesion systems, for example, use nanobody-antigen interactions to stabilize mating pairs and enhance DNA delivery to target recipients, especially under liquid mating conditions where contact formation can limit transfer (20,21). Eex provides a complementary means of controlling recipient selectivity by suppressing plasmid acquisition in cells expressing the corresponding exclusion system. In our experiments, Eex-mediated selectivity was maintained during surface mating, including at increased cell densities and over prolonged mating periods (**Fig. 3**). These findings show that recipient-side exclusion remains effective under conditions that can favor repeated donor– recipient encounters. The complementary roles of adhesion and exclusion suggest that the two strategies could, in principle, be combined. Synthetic adhesion could promote preferential donor contact with target recipients, whereas Eex could suppress plasmid acquisition by non-target recipients engineered to express the cognate exclusion system. Such a combined approach could improve delivery specificity within assembled bacterial communities.

The widespread occurrence of exclusion systems among conjugative plasmids suggests opportunities to extend this strategy to additional conjugation systems. Entry exclusion has been proposed to be an essential feature of conjugative plasmid biology (16). Its occurrence across diverse plasmid groups suggests that recipient-side exclusion is a broadly distributed mechanism for controlling conjugative DNA transfer. Eex modules from different plasmids may therefore provide additional tools for selective DNA delivery, although their functionality and specificity in heterologous hosts will require experimental validation. In this study, we used the RP4/IncP-1 system as a model because of its broad recipient range. RP4-derived Eex reduced conjugative transfer not only in *E. coli* but also in the environmental bacteria *P. putida* KT2440 and *S. japonicum* UT26 (**Figs. 1A and 5A, B**). Moreover, in a three-species recipient mixture containing these bacteria, Eex expression selectively reduced plasmid acquisition in the species expressing the exclusion module, while the other species remained permissive (**Fig. 5C**). These findings demonstrate that RP4-derived Eex can control recipient selectivity across the tested bacterial hosts and within an assembled multispecies community.

Another important feature of Eex-based control is its potential modularity. Exclusion systems generally discriminate between cognate and non-cognate conjugation machineries, suggesting that different exclusion modules could provide distinct recipient-side gates for DNA delivery (17). We demonstrated this principle by combining RP4-type and F-derived conjugation systems with their corresponding exclusion modules (**Fig. 4A**, **B**). When both donor populations were present in the same mating mixture, each plasmid was preferentially transferred to recipients lacking its cognate exclusion system (**Fig. 4C**). This result shows that recipient selectivity can be maintained during simultaneous DNA delivery through two distinct conjugation systems. In principle, extending this approach to additional compatible conjugation–exclusion pairs could enable selective delivery of multiple genetic cargoes to defined recipient populations within a synthetic community. Such expansion would require assessing cross-exclusion and interactions among the transfer systems. These findings provide a basis for developing multiplexed conjugative DNA delivery tools for bacterial consortia.

Eex-based selectivity is particularly well suited to defined synthetic communities, in which recipient permissiveness can be specified before community assembly. By assigning exclusion modules to selected members, this approach allows subsequent DNA delivery to be directed within an established community. This design requires prior engineering of recipient populations, and its application to unmodified natural communities remains to be established. Although exclusion was not absolute, the substantial reductions in transfer observed here provided sufficient selectivity to direct plasmid delivery and, when coupled with selection, alter recipient population composition (**Fig. 6**). Applications requiring lower levels of non-target transfer could build on this approach through optimization of exclusion-module expression or combination with complementary targeting mechanisms.

The variation in baseline transfer frequencies among recipient species **(Fig. 5)** further highlights the importance of matching conjugation and exclusion systems to the intended hosts. Conjugative host range is shaped by multiple factors, including plasmid-encoded transfer machineries, mating-pair stabilization factors, and recipient-side determinants. Understanding how these factors interact with exclusion specificity could guide the selection of compatible donor–recipient combinations and support the development of additional0 delivery routes. Our results with RP4- and F-derived systems provide a foundation for extending this strategy toward programmable DNA delivery across a wider range of synthetic bacterial communities.

## SUPPLEMENTARY MATERIALS

**Figure S1. Expression of trbK alone modestly reduces RP4-mediated conjugative transfer.**

**Figure S2. Growth of E. coli MG1655 carrying arabinose-inducible entry exclusion genes.**

**Figure S3. Effect of arabinose induction on RP4-mediated entry exclusion.**

**Figure S4. Eex-mediated selectivity is maintained during mobilizable plasmid transfer. Table S1. Primers used in this study**

**Supplementary Data S1**

## ACKNOWLEDGEMENTS

We are grateful to Ms. Airi Tatesawa for her technical assistance and support during the experimental work.

## AUTHOR CONTRIBUTIONS

Kouhei Kishida: Conceptualization, Investigation, Formal analysis, Methodology, Validation, Writing - original draft. Natsumi Ogawa-Kishida: Investigation, Methodology, Writing – review and editing. Leonardo Stari: Writing – review and editing. Yoshiyuki Ohtsubo: Writing – review. Yuji Nagata: Writing – review.

## DATA AVAILABILITY

The numerical source data underlying the figures are provided in Supplementary Data S1. Raw microscopy images and source data underlying this study are available in Zenodo at https://doi.org/10.5281/zenodo.22820958

## Supporting Information

**Figure S1.**
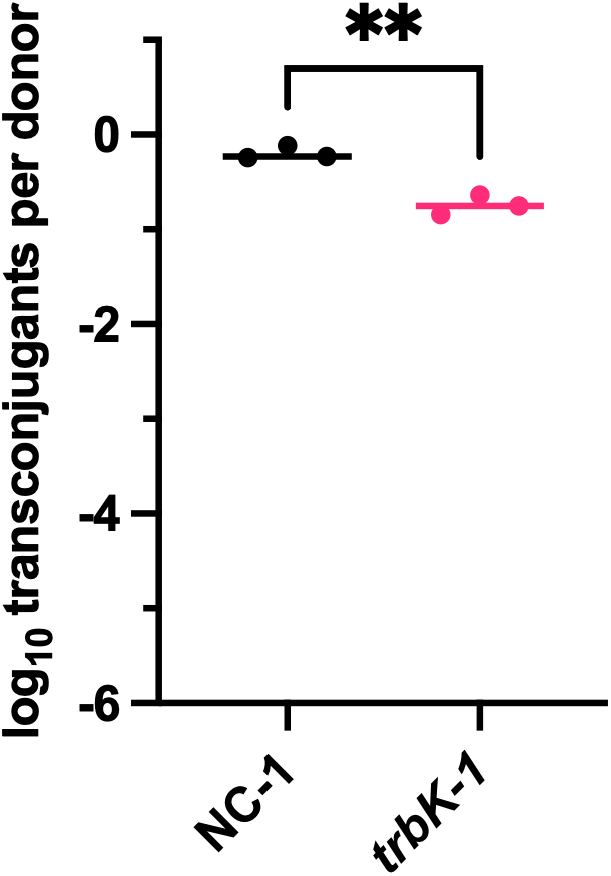
Expression of *trbK* alone modestly reduces RP4-mediated conjugative transfer. Transfer frequencies from the RP4 donor to the control recipient NC-1 and the *trbK*-expressing recipient trbK-1 were calculated as the number of transconjugants per donor. Each dot represents an independent biological replicate (n = 3), and horizontal lines indicate the means. Statistical significance was assessed using an unpaired two-tailed t-test on log₁₀-transformed transfer frequencies. **P < 0.01.

**Figure S2.**
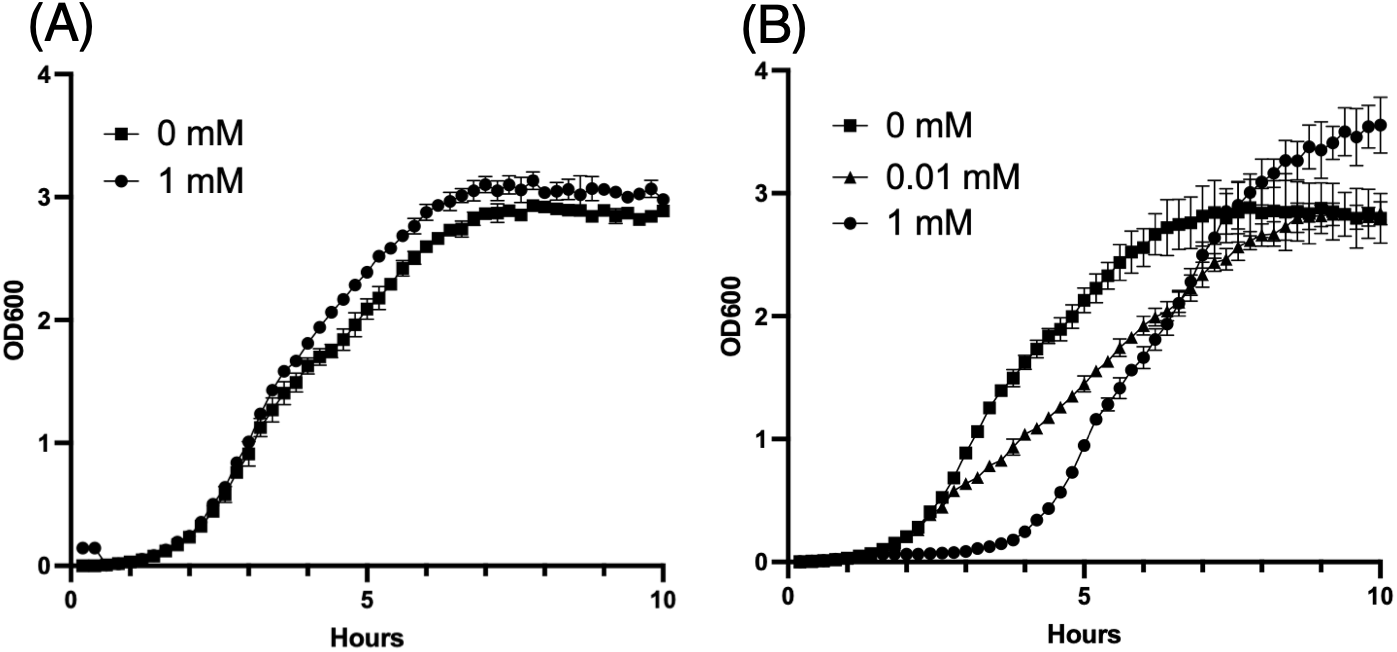
Growth of E. coli MG1655 carrying arabinose-inducible entry exclusion genes. Growth curves of *E. coli* MG1655 carrying (A) *trbJ*–*trbK* or (B) *traS*–*traT* under the control of an arabinose-inducible promoter. Overnight cultures were inoculated at 1% (v/v) into LB medium supplemented with the indicated concentrations of L-arabinose and cultured at 37°C. OD₆₀₀ was measured at 12-min intervals using a biophotorecorder (TVS062CA; ADVANTEC), with the initial reading used as the baseline. Data represent the mean ± SD of three independent biological replicates.

**Figure S3.**
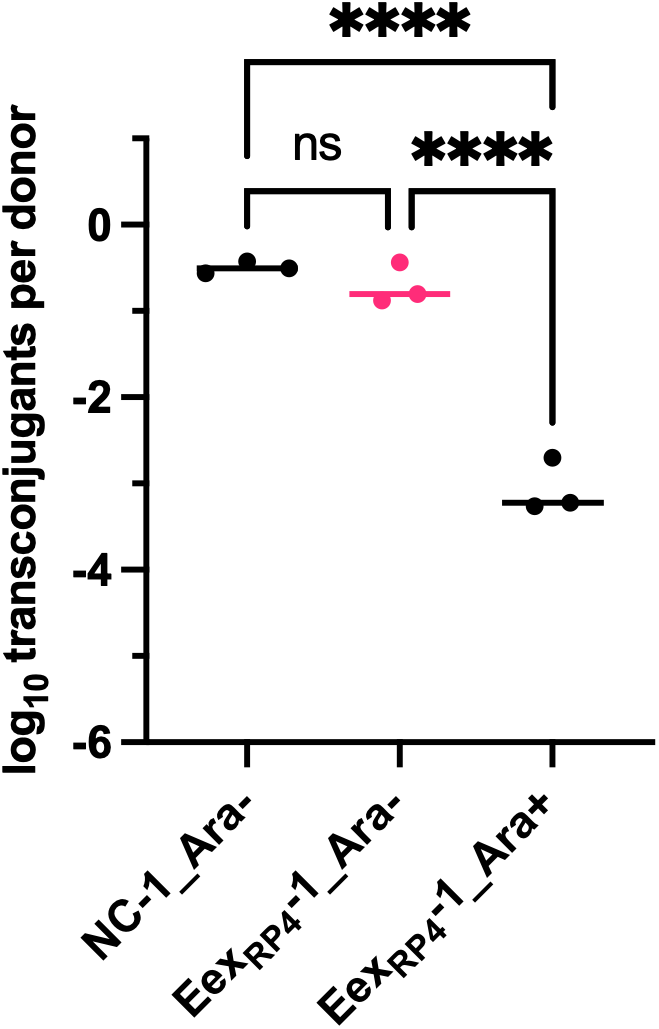
Effect of arabinose induction on RP4-mediated entry exclusion. Conjugative transfer to the control recipient NC-1 in the absence of L-arabinose and to the Eex_RP4_-1 recipient in the absence or presence of L-arabinose. Transfer frequencies are expressed as log₁₀ transconjugants per donor. Each dot represents an independent biological replicate (n = 3), and horizontal lines indicate means of the log₁₀-transformed transfer frequencies. Statistical significance was assessed using ordinary one-way ANOVA followed by Tukey’s multiple-comparisons test on log₁₀-transformed transfer frequencies. ns, not significant; ****P < 0.0001.

**Figure S4.**
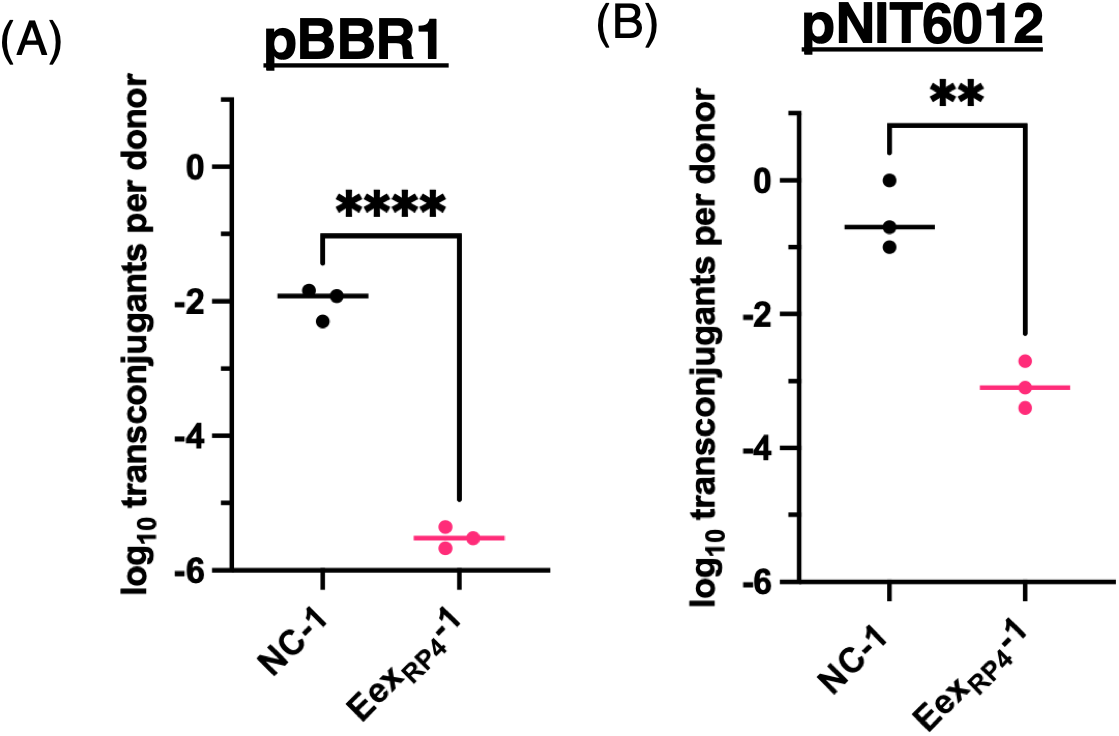
Eex-mediated selectivity is maintained during mobilizable plasmid transfer. **(A)** Quantification of triparental transfer of a mobilizable pBBR1-MCS2 plasmid. A helper donor carrying an RP4-type plasmid, a cargo donor carrying pBBR1-MCS2, and two recipient strains were mixed in the same mating experiment. **(B)** Quantification of pNIT6012 transfer from an *E. coli* S17-1 donor. The S17-1 donor carried pNIT6012, and transfer into control and Eex_RP4_ recipients was quantified in the same recipient mixture. Conjugation frequencies were calculated as transconjugants per donor CFU and are shown on a log_10_ scale. Black dots indicate NC recipients, and magenta dots indicate Eex_RP4_ recipients. Dots represent biological replicates, and horizontal bars indicate the mean. Statistical significance was determined by an unpaired two-tailed t-test. **P < 0.01; ***P < 0.001; ****P < 0.0001.

**Table S1.**
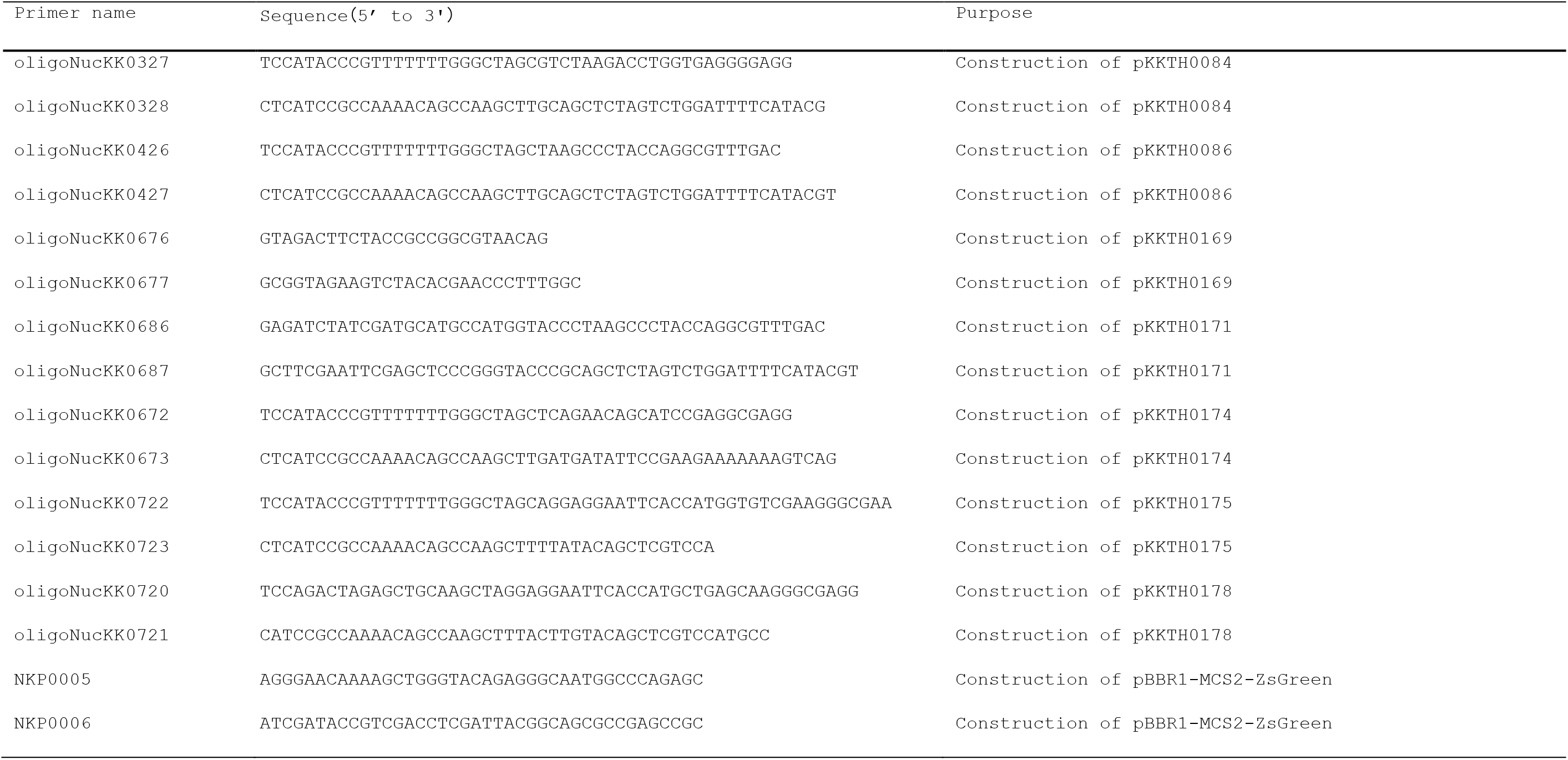
Primers used in this study.

